# Functional divergence of the Arg/N-degron pathway between the crop *Brassica rapa* and the model plant *Arabidopsis thaliana*

**DOI:** 10.64898/2025.12.04.692320

**Authors:** Brian C. Mooney, Pablo Garcia, Shreenivas Kumar Singh, Emmanuelle Graciet

## Abstract

The ubiquitin-dependent Arg/N-degron pathway relates the stability of a substrate protein to the nature of its N-terminal amino acid residue or its biochemical modifications, with some N-terminal residues being recognized by specific E3 ubiquitin ligases, resulting in the ubiquitylation and degradation of the substrate protein. Work in the model plant *Arabidopsis thaliana* has shown that the Arg/N-degron pathway is a key regulator of plant responses to hypoxia, which can be either physiological or a stress in the context of waterlogging or submergence. The role of the Arg/N-degron pathway in hypoxia response is mediated *via* the oxygen-dependent degradation of group VII ETHYLENE RESPONSE FACTOR (ERFVII) transcription factors, which act as the master regulators of the hypoxia response program in plants. Analysis of Arabidopsis mutants for different enzymatic components of the Arg/N-degron pathway has also revealed its roles in the regulation of responses to other abiotic stresses (e.g. salt stress), as well as to pathogens. Although much has been learned from studies in Arabidopsis about the functions of the Arg/N-degron pathway, very little is known about this pathway in crops, including in Brassica crops such as oilseed rape, cabbage or turnip. To determine functional similarities and divergence of the Arg/N-degron pathway between Arabidopsis and Brassica crops, we isolated and characterized the first Arg/N-degron pathway mutants in *Brassica rapa* (turnip, pak choi), a diploid Brassica crop closely related to oilseed rape. We focused on two enzymatic components, namely the arginine-transferases (*ATE*s) and the E3 ubiquitin ligase *PROTEOLYSIS6* (*PRT6*). Our results show both similarities and divergence of function for these Arg/N-degron pathway components in *B. rapa* compared to Arabidopsis. Specifically, *ATE* mutants in *B. rapa* arrest their development at the seedling stage, which contrasts with the mild phenotypic defects of the equivalent Arabidopsis mutants. Double mutant lines for two of the three *PRT6* genes in *B. rapa* indicated a constitutive activation of hypoxia response genes at the transcriptional level, as shown in the single *prt6* mutant in Arabidopsis. However, contrary to Arabidopsis, the *B. rapa* double mutants were more sensitive to waterlogging and hypoxia, and did not show differential response to salt stress or to biotic stress compared to the wild type. The functional divergence identified likely reflects variability in each species in the substrate repertoire and/or in the regulation of pathways or targets downstream of Arg/N-degron pathway substrates. Such differences could be driven by direct selective pressures at N-termini (e.g. gain or loss of a destabilizing N-terminal residue), or by species-specific proteases that may generate destabilizing neo-N-termini after cleavage. These similarities and differences highlight the difficulties in translating research findings from Arabidopsis to crops, even within the same plant family (Brassicaceae) and highlight the need to study pathways in crops.

## Introduction

The ubiquitin/proteasome system plays essential roles in the regulation of plant responses to biotic and abiotic stresses. In plants, the ubiquitin-dependent N-degron pathway in particular functions as a key regulator of plant responses to low oxygen conditions (hypoxia) (reviewed in (Dissmeyer, 2019; Varshavsky, 2019)), which can be caused by environmental conditions such as flooding (including waterlogging or submergence) (Loreti and Perata, 2020; Weits et al., 2021). Work in the model plant *Arabidopsis thaliana* has uncovered the biochemical mechanisms underpinning the role of the N-degron pathway in mediating response to hypoxia (Gibbs et al., 2011; Licausi et al., 2011; Weits et al., 2014), but much less is known about the roles of this pathway in other plants, especially crops. The N-degron pathways relate the stability of a protein to the identity of its N-terminal residue or its post-translational modifications (reviewed in (Dissmeyer, 2019; Varshavsky, 2019)). The so-called Arg/N-degron pathway includes a number of enzymatic components that act (sequentially) to modify a substrate’s N-terminal residue and bind so-called N-terminal destabilizing residues that can act as a degradation signal (degron) (Figure 1A) (Potuschak et al., 1998; Stary et al., 2003; Garzon et al., 2007; Graciet et al., 2009; Graciet et al., 2010). Several *bona fide* Arg/N-degron pathway substrates have been identified in Arabidopsis (Gibbs et al., 2011; Licausi et al., 2011; Gibbs et al., 2018; Goslin et al., 2019; Weits et al., 2019; Labandera et al., 2021), some of which are degraded in an oxygen-dependent manner, such as for example LITTLE ZIPPER 2 (ZPR2) and VERNALIZATION2 (VRN2), which regulate developmental processes in the context of physiological hypoxic niches (Gibbs et al., 2018; Weits et al., 2019; Labandera et al., 2021).

**Figure 1:**
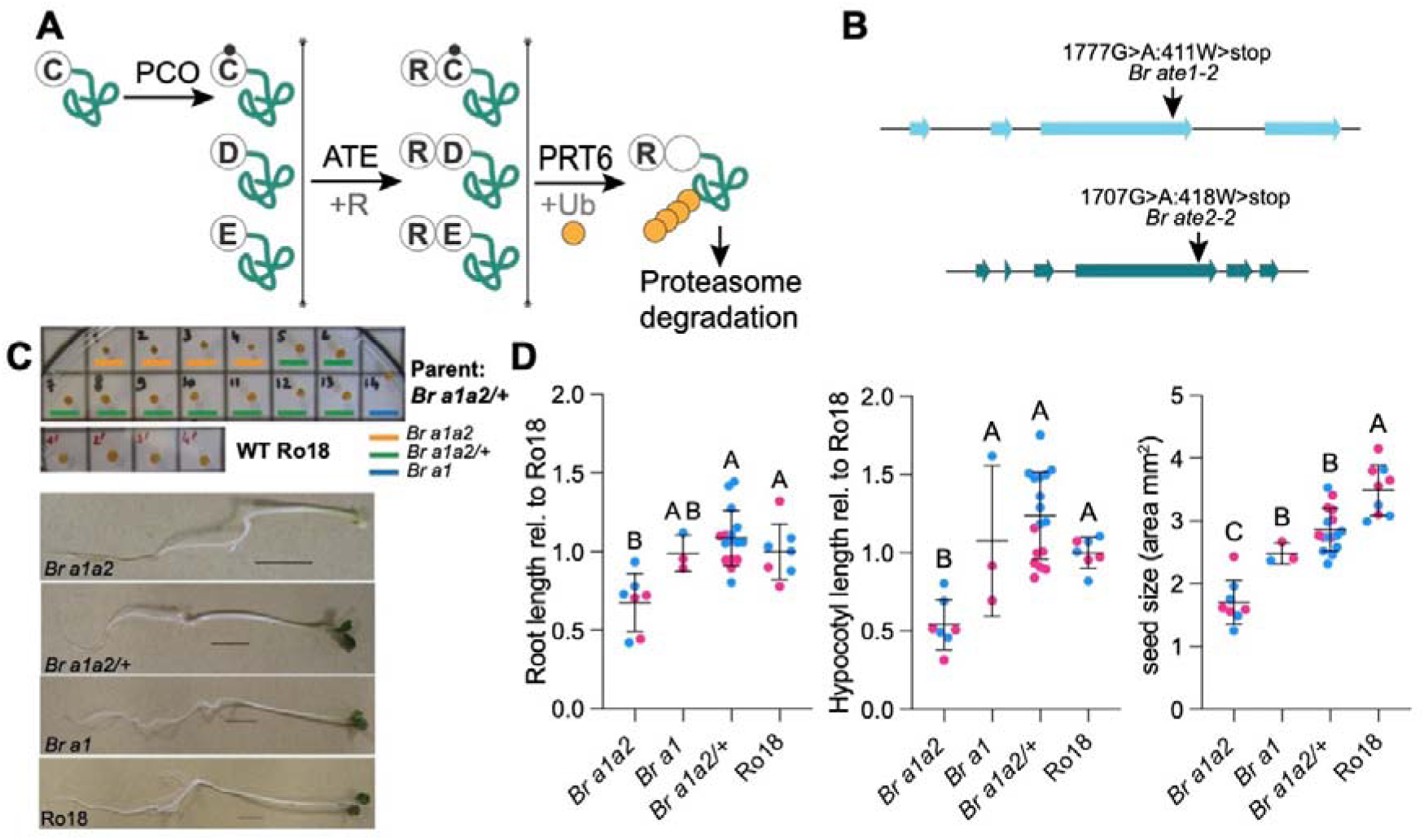
Phenotype of a *Br a1a2* mutant. **(A)** Overview of the Arg/N-degron pathway and its enzymatic components. PLANT CYSTEINE OXIDASE (PCO) are oxygen-dependent enzymes that oxidize the N-terminal Cys (C) residue of proteins (Weits et al., 2014; White et al., 2017). Their substrates can then be arginylated by ATE enzymes, which also conjugate Arg (R) to substrates with N-terminal Asp (D) and Glu (E) (Graciet et al., 2009; Graciet et al., 2010). Arginylation is followed by recognition and ubiquitylation mediated by the E3 ubiquitin ligase PROTEOLYSIS 6 (PRT6), and degradation by the proteasome (Garzon et al., 2007). PRT6 also recognizes proteins that start with other positively charged residues such as lysine (not depicted) (Garzon et al., 2007). **(B)** Gene structure of *Br ATE1* and *Br ATE2* and position of the point mutations in *Br ate1-2* (*Br a1*) and *Br ate2-2* (*Br a2*). Base pair numbering starts at the ATG of the genomic locus. Amino acid substitutions and their position in the protein are indicated. W: tryptophan. **(C)** Seed and seedling phenotype in the offspring population of a *Br a1a2*/+ parent compared to the Ro18 wild type. *Br a1a2*: double homozygous mutant; *Br a1a2/+:* homozygous mutant for *Br ATE1* and heterozygous mutant for *Br ATE2*; *Br a1*: single homozygous mutant for *Br ATE1*. Seedlings were grown under continuous light (20°C) on 0.5xMS agar medium supplemented with 1% (w/v) sucrose. Pictures were taken after 4 days of growth. Scale bar = 1 cm. **(D)** Quantitation of the phenotypes shown in (C), including seed size area, root length relative to that of the average Ro18, hypocotyl length relative to the average Ro18. Mean and standard deviations are shown. Statistical analysis: ANOVA and Tukey *post hoc* test; significant differences: p-value < 0.05. Pink symbols: data obtained from offspring of parent #5 (*Br a1a2*/+); blue symbols: data from offspring of parent #8 (*Br a1a2*/+). Each symbol represents individual seeds/seedlings from a given biological replicate. Data from 2 biological replicates.

The best known substrates of the Arg/N-degron pathway are a set of five group VII ETHYLENE RESPONSE FACTOR transcription factors (collectively noted ERFVIIs) that act as the master regulators of the transcriptional hypoxia response program (Mustroph et al., 2009; Gibbs et al., 2011; Licausi et al., 2011; Reynoso et al., 2019). Their Arg/N-degron-dependent degradation renders this pathway a key regulator of hypoxia responses and a component of the oxygen sensing mechanisms in plants. Arabidopsis *prt6* (*proteolysis 6*) mutants, which are affected for the E3 ubiquitin ligase that ubiquitylates the ERFVIIs, exhibit increased tolerance to waterlogging and to hypoxia due to the constitutive accumulation of the ERFVIIs (Gibbs et al., 2011). Notably, Arabidopsis *prt6* mutants also exhibit increased tolerance to other abiotic stresses such as high salt and drought (Vicente et al., 2017). This phenotype is conserved in barley (*Hordeum vulgare*) plants expressing an RNAi construct targeting *Hv PRT6*, suggesting that Arg/N-degron pathway components could be targets of interest to improve crop tolerance to waterlogging and potentially to other abiotic stresses (Mendiondo et al., 2016; Vicente et al., 2017), including osmotic stress (Papdi et al., 2015). In line with this idea, the overexpression of ERFVII homologs in crop species such as maize (Yu et al., 2019) and wheat (Wei et al., 2019) results in improved tolerance to waterlogging. Overexpression in Arabidopsis of the *RAP2.2* ERFVII transcription factor also led to improved resistance to the fungal necrotrophic pathogen *Botrytis cinerea*, suggesting that the function of these ERFVIIs may extend to the regulation of plant defenses against pathogens (Zhao et al., 2012), perhaps due to the formation of local hypoxic niches at the site of *B. cinerea* infection (Valeri et al., 2021). Similarly, overexpression in Arabidopsis of the barley *RAP2.2* homolog (noted *Hv RAF*) also resulted in increased resistance to the bacterial pathogen *Ralstonia solanacearum* (Jung et al., 2007). Notably, Arabidopsis mutants for Arg/N-degron pathway components also showed differential defense responses to a range pathogens (de Marchi et al., 2016; Gravot et al., 2016; Vicente et al., 2019). Despite this knowledge and potential for applications of agronomic interest, thus far, very little is known about the roles of the Arg/N-degron pathway in crops and the properties of crop plants deficient for some of its components.

As a member of the *Brassicaceae* family, Arabidopsis is related to a range of economically important crop species, including *Brassica napus* (oilseed rape), *Brassica oleracea* (cabbage and broccoli) and *Brassica rapa* (e.g. pak choi, turnip and some oil varieties). *B. rapa* may be considered as a more attractive representative model species for Brassica crops because (i) its diploid genome is less complex than that of the allotetraploid *B. napus*; (ii) the *B. rapa* Chiifu-401-42 genome was sequenced over ten years ago (Wang et al., 2011); (iii) a TILLING (Targeting Induced Local Lesions In Genomes) collection of EMS mutants is available (Stephenson et al., 2010); and (iv) transient expression methods have been optimized (Mooney and Graciet, 2020). Here, we sought to establish *B. rapa* as a model system to investigate the functions of the Arg/N-degron pathway in Brassicaceae crops. Our earlier work revealed that the enzymatic components of the Arg/N-degron pathway identified in Arabidopsis are conserved in *B. rapa*, alongside additional homologs (Mooney and Graciet, 2020). For example, PRT6 is encoded by a single gene in Arabidopsis, but by three genes in *B. rapa*. However, similarly to Arabidopsis, *B. rapa* codes for two Arg-transferases. For the first time, we isolated and characterized *B. rapa* mutants for Arg/N-degron pathway components (specifically, Arg-transferases and PRT6) from the Ro18 TILLING population (Stephenson et al., 2010), and show that they have both similar and divergent phenotypes to their Arabidopsis counterparts. These differences could be relevant to the discovery of new functions of Arg/N-degron pathway in plants and to efforts to translate findings from Arabidopsis to Brassica crops.

## Materials and Methods

### Plant growth media and conditions

*B. rapa* plants were grown on a sterilized soil mixture containing a 5:3:2 ratio of compost, vermiculite and perlite. For experiments involving seedlings, *B. rapa* was grown in Petri-dishes or sterile plastic cups containing 0.5x Murashige and Skoog (MS) medium (pH 5.7) with 6 g/L agar and 0.5% (w/v) sucrose unless stated otherwise in the figure legends. Trays or plates were incubated in the dark at 4°C for 3 days prior to transfer to growth rooms. Plants were grown either in continuous light or in short-day conditions (8 hrs light/16hrs dark), as specified below.

### *B. rapa* lines isolated

*B. rapa* subsp. trilocularis (Yellow Sarson) genotype Ro18 was used. Arg/N-degron mutant lines were isolated from a TILLING population derived from Ro18 EMS mutagenesis (Stephenson et al., 2010) (Supplementary Table S1). In this collection, about one mutation per 60 kb is expected (Stephenson et al., 2010). Seeds were obtained from RevGen UK (John Innes Centre, Norwich). The *Br ate1-2, Br prt6.2-12* and *Br prt6.3-1* lines were backcrossed twice to the wild-type Ro18 parent and *Br ate2-2* was backcrossed once prior to crossing to *Br ate1-2*. After crossing, F1 heterozygous plants for both genes were allowed to self-fertilize in order to obtain segregating populations that could be used to screen for mutant combinations.

### Genotyping of Arg/N-degron mutant *B. rapa* lines

Genomic DNA was extracted as in (Edwards et al., 1991). The *Br ate1-2, Br ate2-2* and *Br prt6.3-1* single nucleotide polymorphisms (SNPs) were genotyped using Sanger sequencing following amplification of the relevant genomic region by PCR using oligonucleotides BM28/BM29, BM30/BM31 and BM36/BM37, respectively (see Supplementary Table S2 for oligonucleotide sequences). A dCAPS assay was used to identify the *Br prt6.2-12* SNP: a nested PCR was carried out using the BM93/BM94 oligonucleotides for the external PCR and BM97/BM98 for the internal PCR, followed by digestion with *Bcl*I for 6 hrs at 55°C (wild type: 219 bp; *Br prt6.2-12*: 199 bp and 20 bp). Presence of the mutation was confirmed by Sanger sequencing after PCR with BM34/BM35.

### Reverse transcription quantitative PCR (RT-qPCR)

Total RNA was extracted using the Spectrum Plant Total RNA Kit (Merck). Reverse transcription reactions were set up using 100-1000 ng of isolated total RNA using RevertAid Reverse Transcriptase (Thermo Fisher), RiboLock RNase inhibitor (Thermo Fisher) and oligo(dT)18. qPCR reaction mixtures were set up in 96-well plates (Roche) with 1 µL of cDNA mixed with 1 µL of a primer pair mixture (1 μM final concentration each) and 5 µL 2X SYBR green master mix (Roche), with nuclease-free water added to a final volume of 10 μL *per* well. qPCR reactions were carried out in a LightCycler 480 instrument (Roche). The second derivative maximum method was used to determine crossing point (Cp) values. Gene expression was calculated relative to a reference gene with the comparative Ct method (Cp_reference gene_ – Cp_gene of interest_ = deltaCp). Assuming a PCR efficiency value of 2, relative expression was calculated as 2^deltaCp^. *Br GAPDH* (*GLYCERALDEHYDE 3-PHOSPHATE DEHYDROGENASE;* Bra016729) was used as a reference gene for RT-qPCRs in *B. rapa* (Procko et al., 2014). Oligonucleotides used for qPCR are listed in Supplementary Table S2.

### Agroinfiltration and transient expression in *B. rapa*

This procedure was carried out as described in (Mooney and Graciet, 2020). *A. tumefaciens* C58 pGV2260 (McBride and Summerfelt, 1990) transformed with the indicated Arg/N-degron pathway reporters (Graciet et al., 2010) or a pML-BART empty vector were grown for 3-4 days at 28°C on LB agar supplemented with 50 mg/L rifampicin, 100 mg/L ampicillin and 100 mg/L spectinomycin. After 3-4 days growth, bacteria were suspended from plates in 2 mL infiltration medium (10 mM MES pH5.5, 10 mM MgCl_2_, 150 µM acetosyringone) and diluted to OD_600_ of 0.75. Four to five week-old *B. rapa* plants were covered with plastic lids overnight prior to infiltration. A ∼2 cm diameter area was marked on the abaxial side of the first and second true leaves. Using a blunt 1 mL syringe, the bacterial suspension was infiltrated into the marked areas. Excess liquid was removed with tissue paper and plants were returned to the growth room. Tissue was harvested 3 days post agroinfiltration for protein or RNA extraction.

### LUC and GUS activity assays with Arg/N-degron pathway reporter constructs

All Arg/N-degron pathway reporter constructs used in this study have been previously published in (Worley et al., 1998; Graciet et al., 2010). The same protocol as in (Mooney and Graciet, 2020) was used. Briefly, proteins were extracted from frozen ground tissue using 1X Luciferase Cell Culture Lysis Reagent (Promega), supplemented with 1 mM phenylmethylsulfonyl fluoride (PMSF) and 1:100 plant Protease Inhibitor Cocktail (Merck). Samples were centrifuged at 12,000 x g for 10 minutes at 4°C to pellet cellular debris. Protein concentration was determined using the Bradford protein assay.

LUC activity was measured as in (Luehrsen et al., 1992; Graciet et al., 2010; Mooney and Graciet, 2020). Briefly, CCLR protein extract (1-2 µL) was added to 100 µL LAR buffer (20 mM tricine, pH7.8, 1.07 mM (MgCO_3_)_4_, Mg(OH)_2_.5H_2_O, 2.67 mM MgSO_4_, 0.1 mM ethylenediaminetetraacetic acid (EDTA), 33.3 mM dithiothreitol (DTT), 270 µM coenzyme A, 470 µM luciferin, 530 µM ATP) in a 96-well plate (Sterilin). Luminescence was measured using a POLARstar Omega microplate reader (BMG LABTECH) for 10 seconds. Luminescence values were then normalized to the relative expression of the *LUC* gene as determined by RT-qPCR from the same tissue (see Supplementary Table S2 for oligonucleotide sequences).

### *B. rapa* flg22 treatment for RNA-Seq

*B. rapa* seedlings were grown in continuous light conditions at 20°C in cups containing 0.5x MS agar supplemented with 0.5% (w/v) sucrose. After 3 days, 4 seedlings *per* genotype *per* treatment were transferred to a well of a 6-well plate containing 6 mL of 0.5x MS supplemented with 0.5% sucrose (liquid medium) and returned to the growth room for incubation overnight with mild shaking. On day 4, seedlings were treated with 1 µM flg22 or an equivalent volume of water (mock treatment) for 1 hr. Seedlings were collected and frozen immediately in liquid nitrogen prior to RNA extraction with Spectrum Plant Total RNA Kit (Merck). Samples from three biological replicates were used.

### RNA-Seq, data processing and functional analysis

RNA integrity was assessed using an Agilent 2100 Bioanalyzer (Agilent). All RNA samples had RNA integrity (RIN) values >7.0. Library preparation and paired-end 100 bp next-generation sequencing was performed by BGI (Hong Kong) using the DNB-seq platform. Data processing was carried out by BGI using the filtering software SOAPnuke (including removal of reads containing the adaptor; removal of reads whose N content is greater than 5%; and removal of low-quality reads). The Hierarchical Indexing for Spliced Alignment of Transcripts (HISAT2) software was then used for mapping clean reads to the *B. rapa* reference genome available at the time of the study (GCF_000309985.2). Differential gene expression was determined using DESeq2. Cut-offs of adjusted p-value < 0.001 and |log_2_ of fold-change| > 1 were applied to determine differentially expressed genes (DEGs). Significant GO category enrichment analyses of DEGs were carried out using ShinyGO 0.85 (Ge et al., 2020) using the STRING.51351 database as reference (STRING v11.5). The following settings were used: FDR cut-off = 0.05, minimum pathway size = 2, maximum pathway size of 5,000, Top 25 GO categories, with removal of redundancy and abbreviation of pathways. As recommended in (Wijesooriya et al., 2022), background for GO analyses with DEGs in the WT#67^flg/m^ dataset, we used the complete set of genes identified in WT#67 when no cut-offs were applied (corresponding to 36,534 genes; see Supplemental Dataset 1). For GO analyses with DEGs in *Br prt6.2/3#68*^flg/m^, the background used corresponded to all the genes identified in *Br prt6.2/3#68* prior to applying cut-offs (36,351 genes; see Supplemental Dataset 2). Overlap between datasets was determined using InteractiVenn (Heberle et al., 2015), and statistical significance of the overlap between datasets was calculated using 2x2 contingency tables and Chi-square tests. Raw and processed data are submitted to NCBI Gene Expression Omnibus under accession number GSE311966.

### Waterlogging stress experiments

Seeds were sown on sterilized soil, and pots were placed in a tray covered with a transparent lid. After 3 days at 4°C, the trays were transferred to the greenhouse (16 hrs light/8hrs dark). After one week of growth, the lids were removed and the plants were grown for 3 weeks, at which point waterlogging treatment was applied by keeping water approximately 1 cm above the soil level. After two weeks of waterlogging, SPAD measurements were taken on 2 different locations (each side of the midvein) of leaf 3 using a Multispeq device (PhotosynQ).

### Hypoxia treatment and chlorophyll content determination

Seeds were surface-sterilised using the vapor-phase sterilization method (Lindsey et al., 2017), and sown in plastic glasses with 0.5xMS agar medium. Seeds were germinated and seedlings were grown in continuous light. Seven-day-old seedlings were treated with hypoxia by placing the cups into anaerojars with an anaerogen sachet (Oxoid) for 16 hrs in the dark. Seedlings were then returned to continuous light conditions. After 24 hrs of recovery, fresh weight of the seedlings was determined and pictures were taken for scoring. Chlorophyll was then extracted following the protocol described in (Sumanta et al., 2014). Scores were defined as: 1 = dead seedling (completely white); 2 = seedling more than 40% yellow or white; 3 = seedling approximately 50% white or yellow; 4 = seedling less than 40% yellow or white; 5 = seedling is completely green. Five biological replicates were performed with 4 seedlings/genotype, condition and replicate.

### Salt stress response assay

Seeds were surface sterilized using the bleach vapor method (Lindsey et al., 2017). Sterilized seeds were sown on Petri dishes with 0.8% water agar and the plates were kept for 24 hrs at 20°C in the dark. Seeds were then transferred to fresh 0.8% water agar plates supplemented with NaCl at concentrations of 100 mM, 150 mM and 200 mM. As a control, seeds were transferred to a fresh 0.8% water agar plate without NaCl. Plates were then kept vertical in continuous light conditions. Seedlings were photographed at 24, 72, and 96 hrs post transfer, and root lengths were measured using ImageJ (Schneider et al., 2012). For each genotype and treatment, five seedlings were used, and three independent biological replicates were performed.

### Inoculation with Sclerotinia sclerotiorum

A *S. sclerotiorum* sclerotium isolated from an oilseed rape field in Ireland was used to propagate *S. sclerotiorum*. Subsequently, sclerotia were cultured on Potato Dextrose Agar (PDA; pH 5.2 - 5.5) and incubated at 21°C for 3 days. Actively growing mycelial plugs were then excised using a sterilized cork borer from the leading edge of the colony and transferred to fresh PDA plates for subculturing. The plate was kept for an additional 48 hrs at 21°C to obtain growing mycelia for inoculation.

For infection assays, fully expanded third leaves of *B. rapa* plants were used. Mycelial agar plugs were excised from the colony margin and placed onto the adaxial surface of the detached leaves. Inoculated leaves were then placed on 90 mm square Petri plates containing 0.8% water agar to maintain humidity. Plates containing inoculated leaves were incubated in short-day conditions at 21°C to facilitate infection. Necrotic lesions on the leaves were photographed 24 hrs post inoculation (hpi). Lesion size was measured using the ImageJ software from images acquired from five biological replicates, each containing eight plants *per* treatment and *per* biological replicate.

### Measurement of apoplastic reactive oxygen species (ROS)

*B. rapa* was grown 20°C for 4 weeks in short-day conditions. Discs (1 cm diameter) were taken from leaves of 4-week-old plants with a cork borer. Leaf discs were then carefully divided into 4 quarters with a razor blade. Each quarter-disc was placed into a separate well of a white Sterilin 96-well plate (ThermoScientific) containing 200 μL dH2O with the abaxial leaf surface facing upwards. The plate was then returned to the growth room for a recovery period of at least 3 hrs. 100X stock solutions of luminol (Merck) (17.7 mg/mL in 200 mM KOH) and horseradish peroxidase (HRP) (Fisher Scientific) (10 mg/mL in dH2O) were prepared fresh. 60 μL of a luminescence solution containing 2.8 μL 100X luminol, 2.8 μL 100X HRP and 54.4 μL dH2O was added to each well. The plate was then transferred to a POLARstar Omega microplate reader (BMG LABTECH) and luminescence was detected for 15 minutes to establish a baseline measurement. During this time, a 1.4 μM stock solution of flg22 was prepared in dH2O. 20 μL of a 1.4 μM stock flg22 solution was added to each well, bringing the total volume to 280 μL, resulting in final concentrations of 100 nM flg22, 1X luminol and 1X HRP. Luminescence was detected every 120 seconds for a 60-minute period after addition of flg22.

### Growth inhibition assays in the presence of flg22

*B. rapa* seedlings were grown in continuous light conditions at 20°C in cups containing 0.5x MS agar supplemented with 0.5% (w/v) sucrose. After 3 days, 3 seedlings per genotype per treatment were transferred to a well of a 6-well plate containing 6 mL of 0.5x MS with 0.5% sucrose (liquid medium) supplemented with 100 nM flg22 or an equivalent volume of deionized water (mock). Seedlings were then grown in this liquid culture with mild shaking in continuous light at 20°C for 7 days, at which point seedlings were weighed.

### Statistical analyses

Statistics tests are presented in the figure legends for each of the relevant panels. All statistical tests were performed using GraphPad Prism.

## Results

### Mutation of Arg-transferases in *B. rapa* causes developmental arrest

One of the best studied Arg/N-degron pathway mutants in Arabidopsis is the *ate1-2 ate2-1* double mutant (noted *a1a2*) which is affected for the two functionally redundant Arg-transferases *AtATE1* and *AtATE2* (Graciet et al., 2009). Similarly to Arabidopsis, *B. rapa* codes for two Arg-transferase homologs, *Br ATE1* and *Br ATE2*. Mutant alleles with a premature stop codon were identified for both genes in a TILLING collection generated in the Ro18 background (Stephenson et al., 2010). Specifically, the *Br ate1-2* mutation substitutes a tryptophan residue at amino acid 411 with a stop codon (TGG -> TAG), while *Br ate2-2* presents a stop codon (TGG -> TGA) instead of a tryptophan at amino acid 418 (Figure 1B). After backcrossing each of these mutant lines with the parental Ro18 genotype to reduce the impact of background mutations, *Br ate1-2*/+ and *Br ate2-2*/+ heterozygous mutant plants were crossed to each other. The F2 population was screened for a *Br ate1-2 ate2-2* double homozygous mutant (noted *Br a1a2*), but no double mutant could be identified. In the F2 population, single homozygous mutant *Br ate1-2* and *Br ate2-2* could be isolated (Supplemental Figure S1), as well as *Br ate1-2/+ ate2-2* (heterozygous for *Br ATE1* and homozygous mutant for *Br ATE2*) and *Br ate1-2 ate2-2/+* (homozygous mutant for *Br ATE1* and heterozygous mutant for *Br ATE2*) plants. When F3 seeds from the *Br ate1-2/+ ate2-2* parent were sown, 20 plants grew, 10 of which were *Br ate1-2* single mutants and 10 were *Br ate1-2/+ ate2-2*. Next, seeds from two individual *Br ate1-2 ate2-2/+* plants were collected and the segregating F3 populations obtained from these two *Br ate1-2 ate2-2*/+ parents were grown on 0.5x MS plates supplemented with 0.5% sucrose. Seedlings displaying under-developed, white cotyledons were consistently identified as double homozygous *Br a1a2* mutants by genotyping and represented about ¼ of the populations (Figure 1C). Other seedlings in the segregating population were either *Br ate1-2* or *Br ate1-2 ate2-2/+*. The *Br a1a2* seedlings had shorter hypocotyls and roots compared to the Ro18 parent, *Br ate1-2* or *Br ate1-2 ate2-2/+* seedlings (Figure 1D). Matching genotype and seed morphology further revealed that seeds of the *Br a1a2* double mutants were smaller than those of genotypes which retained at least one functional copy of *Br ATE2* (Figure 1C and Figure 1D), even though *Br ate1-2 ate2-2/+* seeds were also smaller than those of the Ro18 parental genotype. When left on 0.5x MS medium supplemented with 0.5% sucrose, *Br a1a2* seedlings did not develop beyond the seedling stage. Altogether, these data suggest that, in *B. rapa*, Arg-transferases likely play functionally redundant essential roles in early development. Further characterization of *Br a1a2* plants, both phenotypically and biochemically, was not possible.

### Isolation and characterization of a *B. rapa prt6.2/3* double mutant

A previous BLASTp analysis identified 3 potential homologs of Arabidopsis *PRT6* in *B. rapa* (noted, *Br PRT6.1*, *Br PRT6.2* and *Br PRT6.3*) (Mooney and Graciet, 2020), with *Br PRT6.2* being more highly expressed in wild-type Ro18 seedlings than *Br PRT6.1* and *Br PRT6.3* (Figure 2A). TILLING mutant alleles with premature stop codons for each of these *Br PRT6* homologs were identified (Supplemental Table S1). To avoid an early developmental arrest phenotype similar to that observed in *Br a1a2* mutants, we instead isolated a *Br prt6.2 prt6.3* double mutant (noted *Br prt6.2/3*) that retained the function of the lesser-expressed *Br PRT6.1*. The *Br prt6.2-12* and *Br prt6.3-1* mutant alleles were selected because of the presence of premature stop codons at amino acids 1579 (out of 1986) and 1329 (out of 1968), respectively. These single mutants were back-crossed into the Ro18 parental line, and heterozygous individuals were then crossed to each other to isolate a *Br prt6.2/3* double mutant. Two double homozygous mutant lines (noted #68 and #80) were identified in the F2 population. Within the same segregating F2 population, a line wild-type for both *Br PRT6.2* and *Br PRT6.3* (noted WT#67) was also isolated as an additional ‘wild-type’ control for the presence and potential effects of segregating background mutations that may have been retained despite back-crossing with Ro18. Unlike *Br a1a2* double mutant plants, the *Br prt6.2/3* double mutant lines exhibited a wild-type-like ontogeny.

**Figure 2:**
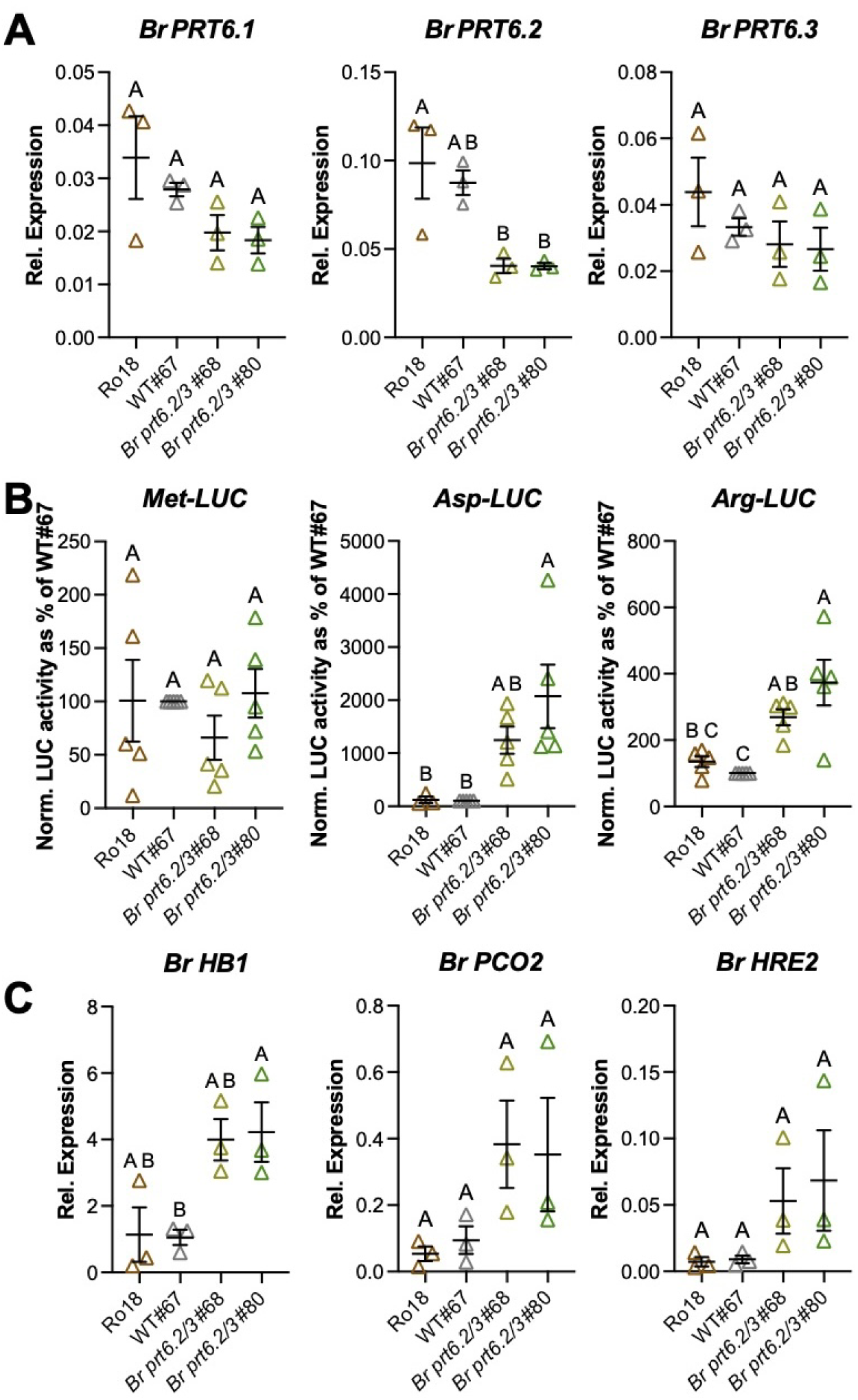
Accumulation of Br PRT6 substrates in *Br prt6.2/3* double mutant lines. **(A)** Expression of *Br PRT6.1*, *Br PRT6.2* and *Br PRT6.3* in 3-day-old seedlings of the wild type and *Br prt6.2/3* mutant. Expression was determined using RT-qPCR. The individual relative expression values to the reference gene (*Br GAPDH*) for 3 biological replicates are shown, together with the mean and standard error of the mean (SEM). Statistical significance (p-value < 0.05) is presented using the CLD format following one-way ANOVA and Tukey’s test. **(B)** Arg/N-degron pathway reporter constructs stability in wild type and *Br prt6.2/3* double mutant lines in transient expression assays. LUC activities normalized for LUC transcript levels were determined and are presented as a percentage of the activity in the WT#67 line. Data represent mean and SEM of 5 biological replicates. Statistical significance (p-value < 0.05) is presented using the CLD format following one-way ANOVA and Tukey’s test. **(C)** Expression of hypoxia response marker genes in 3-day-old wild type and *Br prt6.2/3* seedlings grown on 0.5xMS supplemented with 0.5% sucrose plates. Relative expression levels to the *Br GAPDH* reference gene was determined using RT-qPCR. Mean and SEM of 3 biological replicates are shown. Statistical significance (p-value < 0.05) is presented using the CLD format following one-way ANOVA and Tukey’s test.

Analysis of the expression of *Br PRT6.1*, *Br PRT6.2* and *Br PRT6.3* in the different genetic backgrounds indicated that *Br PRT6.2* mRNA levels were significantly decreased in the *Br prt6.2/3* double mutants, while the mRNA levels of *Br PRT6.1* and *Br PRT6.3* remained similar in the double mutant or in the two wild-type backgrounds used as a control (Figure 2A). Critically, transient expression of Ub-X-LUC Arg/N-degron pathway reporters in leaves of double mutant (*Br prt6.2/3* #68 and #80) and wild-type (Ro18 and WT#67) plants showed significantly increased accumulation of Arg-LUC and Asp-LUC in each of the *Br prt6.2/3* double mutant lines compared to the two wild type lines, while Met-LUC stability was the same irrespective of the genetic background (Figure 2B). To further confirm the disruption of Arg/N-degron pathway function in *Br prt6.2/3*, we tested whether hypoxia response genes were constitutively up-regulated in *Br prt6.2/3* seedlings, as expected from constitutive accumulation of *B. rapa* ERFVII transcription factor homologs. As expected, the expression of *B. rapa* hypoxia response marker genes such as *Br HB1* (Bra001958), *Br PCO2* (Bra025636) and *Br HRE2* (Bra021401) was higher in average (although not always statistically significant) in both *Br prt6.2/3* double mutant lines (#68 and #80) compared to the 2 wild types (Ro18 and #67) (Figure 2C). Altogether, PRT6 activity in *Br prt6.2/3* double mutant plants is sufficiently disrupted to allow accumulation of artificial and natural Arg/N-degron pathway substrates.

### Abiotic stress responses of *Br prt6.2/3* double mutant

One of the most notable phenotypes of *prt6* mutant plants, both in Arabidopsis and in barley, is an increased tolerance to waterlogging and to hypoxia as a result of the constitutive accumulation of ERFVII transcription factors (Gibbs et al., 2011; Mendiondo et al., 2016). The tolerance to waterlogging of *Br prt6.2/3* was therefore examined. Following 15 days of waterlogging of 3-week-old *B. rapa* plants, SPAD measurements were taken as an assessment of relative chlorophyll content. Based on these SPAD values, the waterlogging treatment negatively affected all genotypes, with a stronger negative effect on the two *Br prt6.2/3* mutant lines (Figure 3A). To complement these results, the tolerance of 7-day-old seedlings to hypoxic treatment in the dark for 16 hrs was assessed following a 24-hr recovery period (Figure 3B). The results suggest that the *Br prt6.2/3* double mutant lines are more sensitive to hypoxia than either of the two wild-type controls. To obtain a more accurate assessment of the effect of hypoxia on the seedlings, total chlorophyll levels were determined (Figure 3C). Apart from *Br prt6.2/3* #80 for which no difference between normoxia and hypoxia treatment was apparent, the 3 other genotypes were confirmed to be negatively affected by hypoxia treatment. However, it was not possible to identify relative hypoxia tolerance differences.

**Figure 3:**
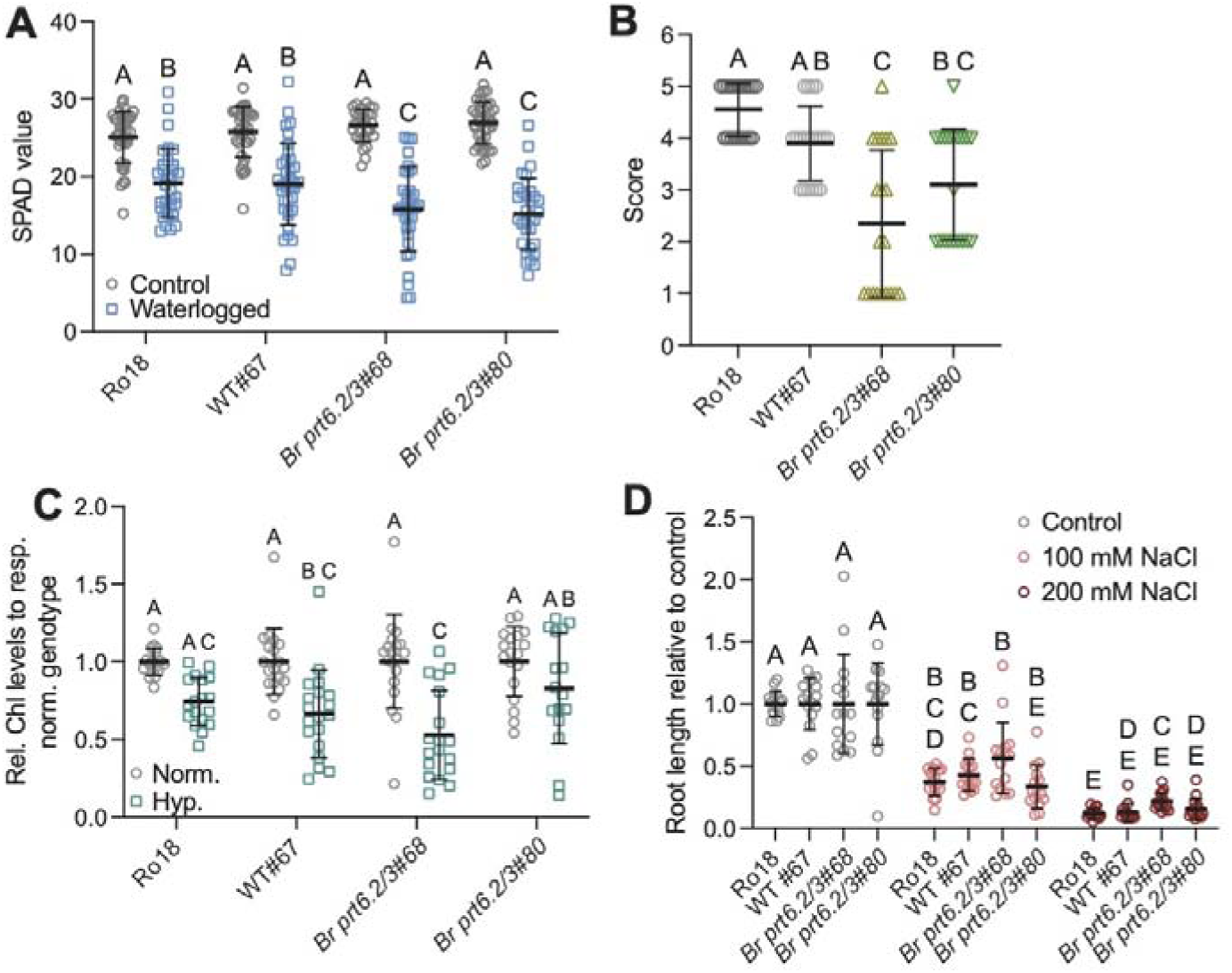
Response of the *Br prt6.2/3* mutant to abiotic stresses. **(A)** SPAD values obtained after a 14-day waterlogging treatment or of control plants kept under normal watering conditions. Mean with standard deviations (SD) of 4 biological replicates (8 plants *per* treatment *per* biological replicate) are shown. Statistical significance was assessed by two-way ANOVA with Sidak’s multiple comparison test. **(B)** Score of seedling health following hypoxia treatment for 16 hrs in the dark, followed by a 24-hr recovery period. Mean scores with SD of 5 biological replicates are shown (4 seedlings/biological replicate). Statistical significance was assessed using one-way ANOVA, followed by Tukey’s test. **(C)** Total chlorophyll (chl; corresponding to chla+chlb) levels relative to those in the same genotype left under normoxic conditions. Mean and SD are shown from 5 biological replicates with 4 seedlings/biological replicate (except replicate 1, which had only 2 seedlings). Two-way ANOVA with Sidak’s multiple comparison test was used to assess statistical significance of differences. **(D)** Root length (in cm) of seedlings grown for 72 hrs with 100 mM or 200 mM NaCl, or mock-treated. For each genotype and treatment, five seedlings were used, and three biological replicates were performed. Two-way ANOVA with Tukey’s multiple comparison test was used to assess statistical significance of differences.

Arabidopsis *prt6* mutants have also been shown to be more tolerant to salt stress (Vicente et al., 2017). Similar tests with *Br prt6.2/3* mutant lines did not detect any differences between the wild-type genotypes and the double mutant lines (Figure 3D). In sum, in contrast to Arabidopsis *prt6* mutant seedlings and plants, *Br prt6.2/3* exhibited increased sensitivity to waterlogging and hypoxia, while salt stress did not reveal any differences between the genotypes.

### Immune and biotic stress responses of *Br prt6.2/3* double mutants

Arabidopsis Arg/N-degron pathway mutants have been shown to be affected for their response to a range of pathogens, albeit with different resistance/susceptibility profiles (de Marchi et al., 2016; Gravot et al., 2016; Vicente et al., 2019). In addition, recent findings have shown connections between the transcriptional response programs to hypoxia and to the model pathogen-associated molecular pattern (PAMP) flg22, which originates from the bacterial flagellin protein and can be used to elicit the first branch of the plant innate immune system, known as pattern-triggered immunity (PTI) (Mooney et al., 2024). To investigate defense-related similarities and differences between Arabidopsis and *B. rapa* Arg/N-degron pathway mutants, innate immune responses of *Br prt6.2/3* double mutants were first compared to those of the two wild-type genotypes using flg22. Gene expression analysis using RT-qPCR indicated that treating 3-day old seedlings with 100 µM flg22 for 1 hr was sufficient to trigger differential expression of two PTI marker genes (*Br MPK3* (Bra038281) and *Br RBOHD* (Bra020724)) in *B. rapa* (Supplemental Fig. S2A). RNA-seq analysis using the same experimental conditions was performed to compare the transcriptomes of the WT#67 and *Br prt6.2/3* lines. Cut-off values of adjusted p-value < 0.001 and |log_2_ of fold-change| > 1 were applied to determine the sets of differentially expressed genes (DEGs) in flg22-treated seedlings compared to mock-treated ones (flg/m) for each genotype. Similar numbers of up- (∼3,900) and down-regulated (∼1,700) genes were identified in WT#67^flg/m^ and in *Br prt6.2/3#68*^flg/m^ (Figure 4A, Supplemental Dataset 1 and Supplemental Dataset 2, respectively). Analysis to identify gene ontology (GO) terms that were enriched in each of these two datasets retrieved expected terms, such as for example ‘defense response’ or ‘response to other organism’ (Figure 4B and Supplemental Dataset 1 and Dataset 2). Most of the top 25 GO categories identified were common to both WT#67^flg/m^ and *Br prt6.2/3#68*^flg/m^ datasets. To further analyze differences in the transcriptional response programs of WT#67 and *Br prt6.2/3#68* to flg22, we determined the overlap between the 2 datasets and identified a statistically significant overlap (p-value < 10^-4^; Chi^2^ test; 4,783 common DEGs) (Figure 4C). In addition, DEGs common to both datasets showed the same directionality of gene expression change and a similar amplitude of up- or down-regulation (Supplemental Fig. S2B).

**Figure 4:**
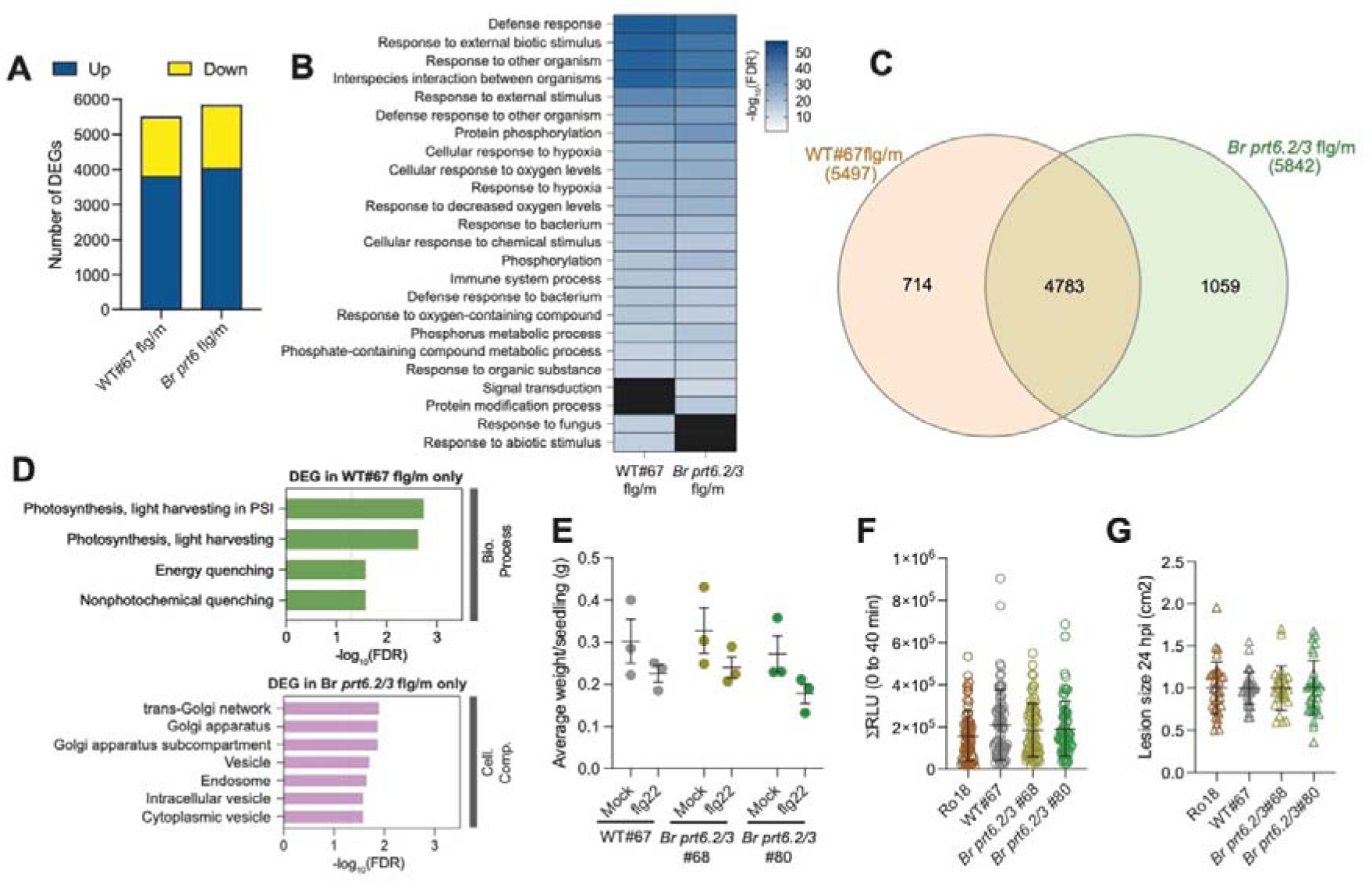
Response of *Br prt6.2/3* mutants to flg22 and *S. sclerotiorum*. **(A)** Number of DEGs in WT#67 and in *Br prt6.2/3*#68 (*Br prt6*) when comparing flg22-treated to mock-treated seedlings (flg/mock) with adj. p <0.001 and |log2FC| > 1.0. **(B)** Top 25 GO Biological Process terms obtained using ShinyGO v0.85 (FDR < 0.05). The GO term ‘Biological proc. involved in interspecies interaction between organisms’ was abbreviated ‘Interspecies interaction between organisms’. **(C)** Overlap and differences between the WT#67^flg/m^ and in *Br prt6.2/3*#68^flg/m^ DEGs. **(D)** GO analysis to determine enrichment for ‘Biological Process’ and also for ‘Cellular Compartment’ amongst the 714 DEGs specific to WT#67^flg/m^ (green) and the 1,059 DEGs in *Br prt6.2/3*#68^flg/m^ (pink) only (FDR < 0.05). PSI: Photosystem I. **(E)** Growth inhibition of 3-day old seedlings co-cultivated with 100 nM flg22 for 7 days. Mean and SEM from 3 biological replicates are shown. No statistically significant differences after 2-way ANOVA. **(F)** ROS production assay. Datapoints represent means of 72 readings (leaf-disc quarters) taken from 18 leaf discs over 3 independent replicates. Error bars indicate SEM. RLU = Relative Light Units. No statistically significant differences after one-way ANOVA. **(G)** Lesion area measured 24 hpi with *S. sclerotiorum*. Mean and SD of 5 biological replicates are shown with 3-8 plants per replicate. No statistically significant differences after one-way ANOVA.

Despite this large overlap, 714 and 1,059 genes were differentially regulated in only one of the 2 genotypes, i.e. in the WT or in the *Br prt6.2/3* mutant, respectively. This suggests some differences in the flg22 response of WT and *Br prt6.2/3* seedlings. GO Biological Process analysis of the 714 DEGs specific to WT#67^flg/m^ revealed an enrichment for GO categories associated with photosynthesis and chloroplast-related processes (Figure 4D), with DEGs in these GO categories being down-regulated in wild-type seedlings in response to flg22 (Supplemental Fig. S2C). A GO analysis for cellular components did not retrieve any statistically significant enrichments. The 1,059 genes differentially expressed in *Br prt6.2/3#68*^flg/m^ only were, in contrast, not found to be enriched for genes associated with specific GO Biological Processes. However, a GO Cellular Component enrichment analysis revealed an over-representation of genes associated with specific compartments such as the ‘Golgi apparatus’ and with the movement of proteins and molecules, including genes associated with ‘intracellular vesicle’ for example. This points to (i) a potential role of Br PRT6 enzymes in the regulation of chloroplast-related processes in wild-type *B. rapa* treated with flg22; and (ii) a potential dysfunction of protein transport in *Br prt6.2/3* seedlings in response to flg22 compared to the wild type.

We next tested the response of the *Br prt6.2/3* mutant to flg22 in a growth inhibition assay that involves 7-day co-cultivation of 3-day-old seedlings with 100 nM flg22, and found that there were not statistically significant differences between the wild-type lines and the mutants (Figure 4E). Similarly, apoplastic ROS production in response to flg22 treatment was not different between the wild type and the *Br prt6.2/3* mutant lines (Figure 4F). Finally, we assessed whether the *Br prt6.2/3* mutants exhibited a different level of resistance to the necrotrophic fungus *Sclerotinia sclerotiorum*, but found no differences in lesion size area between the genotypes tested (Figure 4G). In sum, the flg22 response transcriptional program of *Br prt6.2/3* exhibits differences compared to that of wild-type *B. rapa* seedlings, but this does not translate into defects in other PTI assays or in response to *S. sclerotiorum*.

## Discussion

The Arg/N-degron pathway has been studied in detail in the model plant *Arabidopsis thaliana*, but much less is known about the functions of its enzymatic components in crop species, especially those of the Brassicaceae family such as the diploid *B. rapa*, a close relative of the allotetraploid *B. napus* (oilseed rape). Here, we isolated and characterized the first *B. rapa* Arg/N-degron pathway mutants from a TILLING collection (Stephenson et al., 2010), focusing on two enzymatic components of the Arg/N-degron pathway, the Arg-transferases ATE1/2 and the downstream E3 ubiquitin ligase PRT6 (Figure 1A). Strikingly, the *Br a1a2* double mutant exhibited early developmental arrest, which is in stark contrast to the mild developmental defects of Arabidopsis *ate1 ate2* double mutant plants (i.e. leaf morphology defects, early outgrowth of axillary meristems, phyllotaxis defects and delayed leaf senescence) (Yoshida et al., 2002; Graciet et al., 2009). This phenotype also appears to be stronger than that of *ATE* knock out lines of *Physcomitrella patens*, which exhibited delayed development (Schuessele et al., 2016). Although it remains possible that another co-segregating mutation may contribute to the early developmental arrest phenotype observed, the isolation of single *Br ate1-2* and *Br ate2-2* mutant plants, as well as the use of segregating F3 populations from either *Br ate1-2/+ ate2-2* or *Br ate1-2 ate2-2/+* parents strongly suggests an essential role of Arg-transferases in the regulation of cellular processes at early stages of development, possibly in the shoot apical meristem. In Arabidopsis, the shoot meristem regulator ZPR2 is the only ZPR protein targeted for degradation by the Arg/N-degron pathway in an oxygen dependent manner due to its N-terminal Cys residue (Weits et al., 2019) (ZPR1/3/4 in Arabidopsis do not start with N-terminal Cys). ZPR proteins interact with class III homeodomain leucine zipper (HD-ZIP III) transcription factors, including REVOLUTA (REV), PHABULOSA (PHB) and PHAVOLUTA (PHV) and negatively regulate their activity (Kim et al., 2008; Gruber et al., 2021). Hence plants accumulating ZPR proteins, such as Arg/N-degron pathway mutants, could show similar phenotypic defects as mutants with reduced HD-ZIP III transcription factor activity. BLASTp analysis with Arabidopsis ZPR1/2/3/4 retrieved 11 putative ZPR protein homologs in *B. rapa*, 4 of which start with the Met-Cys sequence, which is characteristic of oxygen-dependent substrates of the Arg/N-degron pathway (the initial Met residue is removed by methionine aminopeptidases, to expose Cys at the N-terminus). This increased number of ZPR Arg/N-degron substrates in *B. rapa* could result in a stronger dependency on the Arg/N-degron pathway to regulate ZPR protein accumulation, which may trigger higher ZPR protein levels in *Br a1a2* than in an Arabidopsis *ate1 ate2* mutant. In turn, this could result in a stronger inhibition of HD-ZIP III in *B. rapa* and a more severe phenotype similar to that of Arabidopsis *rev-6 phb-13* double mutant seedlings which exhibit an early arrest of development (Prigge et al., 2005), a phenotype that is similar to that of *Br a1a2* double mutant plants. An additional, non-mutually exclusive, possibility is that the early developmental arrest of *Br a1a2* mutants is due to the accumulation of *B. rapa-*specific Arg-transferase substrates not present in Arabidopsis and whose removal is normally required for seedling development to continue. Interestingly, the developmental arrest phenotype of *Br a1a2* is reminiscent of the embryonic lethality phenotype of *ate1* mice, in which the only *Mm ATE1* gene is knocked out (Kwon et al., 2002; Brower and Varshavsky, 2009), highlighting the essential roles of Arg-transferases across a broader range of eukaryotes. Our observations with *Br a1a2* also illustrate the difficulties of translating results from a model plant such as Arabidopsis to crops (Roeder et al., 2025; Uauy et al., 2025).

To by-pass the early developmental arrest phenotype of *Br a1a2*, which precluded further investigation of the Arg/N-degron pathway, and to avoid working with segregating populations from *Br ate1-2 ate2-2/+* parents, we isolated a *Br prt6.2/3* double mutant line containing early stop codon mutations in the two (out of 3) most strongly expressed *B. rapa* homologs of Arabidopsis *PRT6*. *Br prt6.2/3* mutants developed similarly to wild-type plants, while showing increased stability of Asp-LUC and Arg-LUC reporters indicating that overall PRT6 activity was sufficiently impaired to allow accumulation of Arg/N-degron pathway substrates. This is in agreement with the constitutive up-regulation of hypoxia-response genes in *Br prt6.2/3*, likely due to the stabilization of the ERFVII transcription factors. Hence, the *Br prt6.2/3* lines could be a suitable tool to study the functions of the Arg/N-degron pathway in *B. rapa*.

Our characterization of *Br prt6.2/3* lines focused on responses to (a)biotic stresses for which differences between Arabidopsis wild-type and Arg/N-degron pathway mutant have been published. Arabidopsis *prt6* mutants are more tolerant to hypoxia stress (Gibbs et al., 2011), as well as to submergence (Riber et al., 2015), although for the latter, contrasting observation were made (Licausi et al., 2011). Here, under the experimental conditions applied, *Br prt6.2/3* plants were more susceptible to waterlogging than the wild type (Figure 3A). Submergence and waterlogging assays yield variable results in general, with many different parameters contributing to differential outcome (e.g. light/dark, humidity, etc…). Here, the observed sensitivity of *Br prt6.2/3* compared to the wild type may reflect an increased susceptibility of *Br prt6.2/3* roots to waterlogging stress, which may affect the ability of the mutants to maintain metabolic pathways and physiological processes. The sensitivity could also result from differences in the size of the root systems, although the latter could not be analyzed due to difficulties to maintain an intact root system after waterlogging. Root growth survival assays are also more difficult to carry out with *B. rapa* due to the larger size of the seedlings/plants and the need to grow them in a vertical manner to allow the roots to grow along the medium. Another possibility is that carbon metabolism and starvation response in the context of waterlogging or hypoxia in *B. rapa* may be different from that of Arabidopsis. However, this is unlikely to be the case, as such differences have not been observed when comparing *B. napus* and Arabidopsis transcriptional responses to hypoxia, as well as sugar levels (Ambros et al., 2022). We sought to test the response of the *Br prt6.2/3* mutant to hypoxia. These tests also suggested that the *Br prt6.2/3* mutant is more susceptible to hypoxia, even though conflicting evidence was found between the two *Br prt6.2/3* lines used when measuring chlorophyll levels. Survival in such assays is often associated with sugar metabolism and the ability to withstand starvation, however, Arabidopsis and *B. napus* do not seem to show significant differences in this respect (Ambros et al., 2022), and a similar situation is likely the case with *B. rapa*. In Arabidopsis, *prt6* mutants have been shown to be more salt stress tolerant (Vicente et al., 2017), but *Br prt6.2/3* mutants behaved similarly to the wild type in our assays. One possibility is that the remaining functional allele, Br PRT6.1, is sufficient to mask a phenotype.

In Arabidopsis, Arg/N-degron pathway mutants differ from wild type in terms of their defense response to a range of pathogens (de Marchi et al., 2016; Gravot et al., 2016; Vicente et al., 2019). We identified transcriptional differences in the response of the *Br prt6.2/3* mutant to the model PAMP flg22, specifically in terms of the regulation of photosynthesis-related genes (which were enriched among wild type only DEGs) and of processes involved in cellular transport (enriched amongst DEGs specific to *Br prt6.2/3*). This suggests a potential role of the Arg/N-degron pathway in the regulation of PTI, despite the lack of other PTI-associated defects in the double mutant. We also tested the response of the *Br prt6.2/3* mutant to the necrotrophic fungal pathogen *S. sclerotinia*, but did not observe differences compared to the wild type. This contrasts with the increased susceptibility of the Arabidopsis *ate1 ate2* mutant (de Marchi et al., 2016), and could be the result of the remaining activity of Br PRT6.1 in the double mutant.

In summary, the results from the Arabidopsis/*B. rapa* comparative analyses reinforce the need to validate knowledge gained in model systems *via* direct experimentation in crop species (Roeder et al., 2025; Uauy et al., 2025). For example, the species-specific functions of the Arg/N-degron pathway identified here suggest a partial divergence of the physiological roles of the Arg/N-degron pathway since the split of Arabidopsis from Brassicas 43 million years ago (Beilstein et al., 2010). As the Arg/N-degron pathway components and structure appear to be well-conserved, this likely reflects variability in each species in the substrate repertoire and/or in the regulation of pathways or targets downstream of Arg/N-degron pathway substrates. Such differences could be driven by direct selective pressures at N-termini (e.g. gain or loss of a destabilizing N-terminal residue), or by species-specific proteases that may generate destabilizing neo-N-termini after cleavage.

## Acknowledgements

This work was funded by grants 13/IA/1870 and 20/FFP-P/8433 from Science Foundation Ireland (now Research Ireland) to EG. BCM was supported by an Irish Research Council PhD scholarship (GOIPG/2017/2). We are grateful to Dr. Ewen Mullins (Teagasc Oak Park) for supplying sclerotia of *Sclerotinia sclerotiorum*, as well as to Prof. Frank Wellmer and Dr. Joseph Beegan for helpful advice regarding the analysis of RNA-seq datasets. The authors confirm that they have no conflict of interest.

## Authors contributions

BCM, PG, SS and EG designed the work, conducted experiments, analyzed data and wrote the manuscript.

## Accession numbers

Raw and processed data are submitted to NCBI Gene Expression Omnibus under accession number GSE311966.

## Notes

### Competing Interest Statement

The authors have declared no competing interest.

### Summary of Updates

- an additional Supplementary Figure S1 with pictures of wild-type, ate1-2 and ate2-2 seedlings to demonstrate that an ate2-2 single mutant is viable. - additional text on results we had obtained from the F3 population from ate1-2/+ ate2-2 parent, which indicates that the developmental arrest of the ate1-2 ate2-2 double mutant is unlikely to be linked to a co-segregating mutation with ate2-2. - a revised version of all GO analyses in the manuscript using a background dataset that contained the genes that we could identify in our RNA-seq experiment when no cut-offs are applied (as opposed to all the genes in the genome). - more cautious conclusions considering the mutation frequency in the TILLING population used in this study.

## References

1. Ambros S, Kotewitsch M, Wittig PR, Bammer B, Mustroph A (2022) Transcriptional Response of Two Brassica napus Cultivars to Short-Term Hypoxia in the Root Zone. Front Plant Sci 13: 897673

2. Beilstein MA, Nagalingum NS, Clements MD, Manchester SR, Mathews S (2010) Dated molecular phylogenies indicate a Miocene origin for Arabidopsis thaliana. Proc Natl Acad Sci U S A 107: 18724–18728

3. Brower CS, Varshavsky A (2009) Ablation of arginylation in the mouse N-end rule pathway: loss of fat, higher metabolic rate, damaged spermatogenesis, and neurological perturbations. PLoS One 4: e7757

4. de Marchi R, Sorel M, Mooney B, Fudal I, Goslin K, Kwasniewska K, Ryan PT, Pfalz M, Kroymann J, Pollmann S, Feechan A, Wellmer F, Rivas S, Graciet E (2016) The N-end rule pathway regulates pathogen responses in plants. Sci Rep 6: 26020

5. Dissmeyer N (2019) Conditional Protein Function via N-Degron Pathway-Mediated Proteostasis in Stress Physiology. Annu Rev Plant Biol 70: 83–117

6. Edwards K, Johnstone C, Thompson C (1991) A simple and rapid method for the preparation of plant genomic DNA for PCR analysis. Nucleic Acids Res 19: 1349

7. Garzon M, Eifler K, Faust A, Scheel H, Hofmann K, Koncz C, Yephremov A, Bachmair A (2007) PRT6/At5g02310 encodes an Arabidopsis ubiquitin ligase of the N-end rule pathway with arginine specificity and is not the CER3 locus. FEBS Lett 581: 3189–3196

8. Ge SX, Jung D, Yao R (2020) ShinyGO: a graphical gene-set enrichment tool for animals and plants. Bioinformatics 36: 2628–2629

9. Gibbs DJ, Lee SC, Isa NM, Gramuglia S, Fukao T, Bassel GW, Correia CS, Corbineau F, Theodoulou FL, Bailey-Serres J, Holdsworth MJ (2011) Homeostatic response to hypoxia is regulated by the N-end rule pathway in plants. Nature 479: 415–418

10. Gibbs DJ, Tedds HM, Labandera AM, Bailey M, White MD, Hartman S, Sprigg C, Mogg SL, Osborne R, Dambire C, Boeckx T, Paling Z, Voesenek L, Flashman E, Holdsworth MJ (2018) Oxygen-dependent proteolysis regulates the stability of angiosperm polycomb repressive complex 2 subunit VERNALIZATION 2. Nat Commun 9: 5438

11. Goslin K, Eschen-Lippold L, Naumann C, Linster E, Sorel M, Klecker M, de Marchi R, Kind A, Wirtz M, Lee J, Dissmeyer N, Graciet E (2019) Differential N-end Rule Degradation of RIN4/NOI Fragments Generated by the AvrRpt2 Effector Protease. Plant Physiol 180: 2272–2289

12. Graciet E, Mesiti F, Wellmer F (2010) Structure and evolutionary conservation of the plant N-end rule pathway. Plant J 61: 741–751

13. Graciet E, Walter F, Maoileidigh DO, Pollmann S, Meyerowitz EM, Varshavsky A, Wellmer F (2009) The N-end rule pathway controls multiple functions during Arabidopsis shoot and leaf development. Proc Natl Acad Sci U S A 106: 13618–13623

14. Gravot A, Richard G, Lime T, Lemarie S, Jubault M, Lariagon C, Lemoine J, Vicente J, Robert-Seilaniantz A, Holdsworth MJ, Manzanares-Dauleux MJ (2016) Hypoxia response in Arabidopsis roots infected by Plasmodiophora brassicae supports the development of clubroot. BMC Plant Biol 16: 251

15. Gruber AV, Kosty M, Jami-Alahmadi Y, Wohlschlegel JA, Long JA (2021) The dynamics of HD-ZIP III - ZPR protein interactions play essential roles in embryogenesis, meristem function and organ development. bioRxiv: 2021.2011.2024.469949

16. Heberle H, Meirelles GV, da Silva FR, Telles GP, Minghim R (2015) InteractiVenn: a web-based tool for the analysis of sets through Venn diagrams. BMC Bioinformatics 16: 169

17. Jung J, Won SY, Suh SC, Kim H, Wing R, Jeong Y, Hwang I, Kim M (2007) The barley ERF-type transcription factor HvRAF confers enhanced pathogen resistance and salt tolerance in Arabidopsis. Planta 225: 575–588

18. Kim YS, Kim SG, Lee M, Lee I, Park HY, Seo PJ, Jung JH, Kwon EJ, Suh SW, Paek KH, Park CM (2008) HD-ZIP III activity is modulated by competitive inhibitors via a feedback loop in Arabidopsis shoot apical meristem development. Plant Cell 20: 920–933

19. Kwon YT, Kashina AS, Davydov IV, Hu RG, An JY, Seo JW, Du F, Varshavsky A (2002) An essential role of N-terminal arginylation in cardiovascular development. Science 297: 96–99

20. Labandera AM, Tedds HM, Bailey M, Sprigg C, Etherington RD, Akintewe O, Kalleechurn G, Holdsworth MJ, Gibbs DJ (2021) The PRT6 N-degron pathway restricts VERNALIZATION 2 to endogenous hypoxic niches to modulate plant development. New Phytol 229: 126–139

21. Licausi F, Kosmacz M, Weits DA, Giuntoli B, Giorgi FM, Voesenek LA, Perata P, van Dongen JT (2011) Oxygen sensing in plants is mediated by an N-end rule pathway for protein destabilization. Nature 479: 419–422

22. Lindsey BE, 3rd, Rivero L, Calhoun CS, Grotewold E, Brkljacic J (2017) Standardized Method for High-throughput Sterilization of Arabidopsis Seeds. J Vis Exp 128: 56587

23. Loreti E, Perata P (2020) The Many Facets of Hypoxia in Plants. Plants (Basel) 9: 745

24. Luehrsen KR, de Wet JR, Walbot V (1992) Transient expression analysis in plants using firefly luciferase reporter gene. Meth Enzymol 216: 397–414

25. McBride KE, Summerfelt KR (1990) Improved binary vectors for Agrobacterium-mediated plant transformation. Plant Mol Biol 14: 269–276

26. Mendiondo GM, Gibbs DJ, Szurman-Zubrzycka M, Korn A, Marquez J, Szarejko I, Maluszynski M, King J, Axcell B, Smart K, Corbineau F, Holdsworth MJ (2016) Enhanced waterlogging tolerance in barley by manipulation of expression of the N-end rule pathway E3 ligase PROTEOLYSIS6. Plant Biotechnol J 14: 40–50

27. Mooney BC, Doorly CM, Mantz M, Garcia P, Huesgen PF, Graciet E (2024) Hypoxia represses pattern-triggered immune responses in Arabidopsis. Plant Physiol 196: 2064–2077

28. Mooney BC, Graciet E (2020) A simple and efficient Agrobacterium-mediated transient expression system to dissect molecular processes in Brassica rapa and Brassica napus. Plant Direct 4: e00237

29. Mustroph A, Zanetti ME, Jang CJ, Holtan HE, Repetti PP, Galbraith DW, Girke T, Bailey-Serres J (2009) Profiling translatomes of discrete cell populations resolves altered cellular priorities during hypoxia in Arabidopsis. Proc Natl Acad Sci U S A 106: 18843–18848

30. Papdi C, Perez-Salamo I, Joseph MP, Giuntoli B, Bogre L, Koncz C, Szabados L (2015) The low oxygen, oxidative and osmotic stress responses synergistically act through the ethylene response factor VII genes RAP2.12, RAP2.2 and RAP2.3. Plant J 82: 772–784

31. Potuschak T, Stary S, Schlogelhofer P, Becker F, Nejinskaia V, Bachmair A (1998) PRT1 of Arabidopsis thaliana encodes a component of the plant N-end rule pathway. Proc Natl Acad Sci U S A 95: 7904–7908

32. Prigge MJ, Otsuga D, Alonso JM, Ecker JR, Drews GN, Clark SE (2005) Class III homeodomain-leucine zipper gene family members have overlapping, antagonistic, and distinct roles in Arabidopsis development. Plant Cell 17: 61–76

33. Procko C, Crenshaw CM, Ljung K, Noel JP, Chory J (2014) Cotyledon-Generated Auxin Is Required for Shade-Induced Hypocotyl Growth in Brassica rapa. Plant Physiol 165: 1285–1301

34. Reynoso MA, Kajala K, Bajic M, West DA, Pauluzzi G, Yao AI, Hatch K, Zumstein K, Woodhouse M, Rodriguez-Medina J, Sinha N, Brady SM, Deal RB, Bailey-Serres J (2019) Evolutionary flexibility in flooding response circuitry in angiosperms. Science 365: 1291–1295

35. Riber W, Muller JT, Visser EJ, Sasidharan R, Voesenek LA, Mustroph A (2015) The greening after extended darkness1 is an N-end rule pathway mutant with high tolerance to submergence and starvation. Plant Physiol 167: 1616–1629

36. Roeder AHK, Bent A, Lovell JT, McKay JK, Bravo A, Medina-Jimenez K, Morimoto KW, Brady SM, Hua L, Hibberd JM, Zhong S, Cardinale F, Visentin I, Lovisolo C, Hannah MA, Webb AAR (2025) Lost in translation: What we have learned from attributes that do not translate from Arabidopsis to other plants. Plant Cell 37: koaf036

37. Schneider CA, Rasband WS, Eliceiri KW (2012) NIH Image to ImageJ: 25 years of image analysis. Nat Methods 9: 671–675

38. Schuessele C, Hoernstein SN, Mueller SJ, Rodriguez-Franco M, Lorenz T, Lang D, Igloi GL, Reski R (2016) Spatio-temporal patterning of arginyl-tRNA protein transferase (ATE) contributes to gametophytic development in a moss. New Phytol 209: 1014–1027

39. Stary S, Yin XJ, Potuschak T, Schlogelhofer P, Nizhynska V, Bachmair A (2003) PRT1 of Arabidopsis is a ubiquitin protein ligase of the plant N-end rule pathway with specificity for aromatic amino-terminal residues. Plant Physiol 133: 1360–1366

40. Stephenson P, Baker D, Girin T, Perez A, Amoah S, King GJ, Ostergaard L (2010) A rich TILLING resource for studying gene function in Brassica rapa. BMC Plant Biol 10: 62

41. Sumanta N, Haque CI, Nishika J, Suprakash R (2014) Spectrophotometric Analysis of Chlorophylls and Carotenoids from Commonly Grown Fern Species by Using Various Extracting Solvents. Research Journal of Chemical Sciences 4: 63–69

42. Uauy C, Nelissen H, Chan RL, Napier JA, Seung D, Liu L, McKim SM (2025) Challenges of translating Arabidopsis insights into crops. Plant Cell 37: koaf059

43. Valeri MC, Novi G, Weits DA, Mensuali A, Perata P, Loreti E (2021) Botrytis cinerea induces local hypoxia in Arabidopsis leaves. New Phytol 229: 173–185

44. Varshavsky A (2019) N-degron and C-degron pathways of protein degradation. Proc Natl Acad Sci U S A 116: 358–366

45. Vicente J, Mendiondo GM, Movahedi M, Peirats-Llobet M, Juan YT, Shen YY, Dambire C, Smart K, Rodriguez PL, Charng YY, Gray JE, Holdsworth MJ (2017) The Cys-Arg/N-End Rule Pathway Is a General Sensor of Abiotic Stress in Flowering Plants. Curr Biol 27: 3183–3190 e3184

46. Vicente J, Mendiondo GM, Pauwels J, Pastor V, Izquierdo Y, Naumann C, Movahedi M, Rooney D, Gibbs DJ, Smart K, Bachmair A, Gray JE, Dissmeyer N, Castresana C, Ray RV, Gevaert K, Holdsworth MJ (2019) Distinct branches of the N-end rule pathway modulate the plant immune response. New Phytol 221: 988–1000

47. Wang X, Wang H, Wang J, Sun R, Wu J, Liu S, Bai Y, Mun JH, Bancroft I, Cheng F, Huang S, Li X, Hua W, Wang J, Wang X, Freeling M, Pires JC, Paterson AH, Chalhoub B, Wang B, Hayward A, Sharpe AG, Park BS, Weisshaar B, Liu B, Li B, Liu B, Tong C, Song C, Duran C, Peng C, Geng C, Koh C, Lin C, Edwards D, Mu D, Shen D, Soumpourou E, Li F, Fraser F, Conant G, Lassalle G, King GJ, Bonnema G, Tang H, Wang H, Belcram H, Zhou H, Hirakawa H, Abe H, Guo H, Wang H, Jin H, Parkin IA, Batley J, Kim JS, Just J, Li J, Xu J, Deng J, Kim JA, Li J, Yu J, Meng J, Wang J, Min J, Poulain J, Wang J, Hatakeyama K, Wu K, Wang L, Fang L, Trick M, Links MG, Zhao M, Jin M, Ramchiary N, Drou N, Berkman PJ, Cai Q, Huang Q, Li R, Tabata S, Cheng S, Zhang S, Zhang S, Huang S, Sato S, Sun S, Kwon SJ, Choi SR, Lee TH, Fan W, Zhao X, Tan X, Xu X, Wang Y, Qiu Y, Yin Y, Li Y, Du Y, Liao Y, Lim Y, Narusaka Y, Wang Y, Wang Z, Li Z, Wang Z, Xiong Z, Zhang Z (2011) The genome of the mesopolyploid crop species Brassica rapa. Nat Genet 43: 1035–1039

48. Wei X, Xu H, Rong W, Ye X, Zhang Z (2019) Constitutive expression of a stabilized transcription factor group VII ethylene response factor enhances waterlogging tolerance in wheat without penalizing grain yield. Plant Cell Environ 42: 1471–1485

49. Weits DA, Giuntoli B, Kosmacz M, Parlanti S, Hubberten HM, Riegler H, Hoefgen R, Perata P, van Dongen JT, Licausi F (2014) Plant cysteine oxidases control the oxygen-dependent branch of the N-end-rule pathway. Nat Commun 5: 3425

50. Weits DA, Kunkowska AB, Kamps NCW, Portz KMS, Packbier NK, Nemec Venza Z, Gaillochet C, Lohmann JU, Pedersen O, van Dongen JT, Licausi F (2019) An apical hypoxic niche sets the pace of shoot meristem activity. Nature 569: 714–717

51. Weits DA, van Dongen JT, Licausi F (2021) Molecular oxygen as a signaling component in plant development. New Phytol 229: 24–35

52. White MD, Klecker M, Hopkinson RJ, Weits DA, Mueller C, Naumann C, O’Neill R, Wickens J, Yang J, Brooks-Bartlett JC, Garman EF, Grossmann TN, Dissmeyer N, Flashman E (2017) Plant cysteine oxidases are dioxygenases that directly enable arginyl transferase-catalysed arginylation of N-end rule targets. Nat Commun 8: 14690

53. Wijesooriya K, Jadaan SA, Perera KL, Kaur T, Ziemann M (2022) Urgent need for consistent standards in functional enrichment analysis. PLoS Comput Biol 18: e1009935

54. Worley CK, Ling R, Callis J (1998) Engineering in vivo instability of firefly luciferase and Escherichia coli beta-glucuronidase in higher plants using recognition elements from the ubiquitin pathway. Plant Mol Biol 37: 337–347

55. Yoshida S, Ito M, Callis J, Nishida I, Watanabe A (2002) A delayed leaf senescence mutant is defective in arginyl-tRNA:protein arginyltransferase, a component of the N-end rule pathway in Arabidopsis. Plant J 32: 129–137

56. Yu F, Liang K, Fang T, Zhao H, Han X, Cai M, Qiu F (2019) A group VII ethylene response factor gene, ZmEREB180, coordinates waterlogging tolerance in maize seedlings. Plant Biotechnol J 17: 2286–2298

57. Zhao Y, Wei T, Yin KQ, Chen Z, Gu H, Qu LJ, Qin G (2012) Arabidopsis RAP2.2 plays an important role in plant resistance to Botrytis cinerea and ethylene responses. New Phytol 195: 450–460

