## Supplement for "Functional divergence of the Arg/N-degron pathway between the crop *Brassica rapa* and the model plant *Arabidopsis thaliana*"

^#^ Equal author contribution

**Supplementary Table S1: List of *B. rapa* lines used and isolated in this study.**

| **Line** | **Description** | **Reference** |
| --- | --- | --- |
| Ro18 | Wild-type Ro18 seeds from John Innes Centre (JIC). | (Stephenson et al., 2010) |
| *Br prt6.2-12* | Contains early stop codon in *Br PRT6.2* (Bra009598). Obtained from the JIC; line #JI40184-A. | This study |
| *Br prt6.3-1* | Contains early stop codon in *Br PRT6.3* (Bra028876). Obtained from the JIC; line #JI32812-B. | This study |
| *Br prt6.2/3 #68* | Double mutant isolated from cross between *Br prt6.2-12* and *Br prt6.3-1*, line #68. | This study |
| *Br prt6.2/3 #80* | Double mutant isolated from cross between *Br prt6.2-12* and *Br prt6.3-1*, line #80. | This study |
| WT #67 | ‘Wild-type’ line isolated from F2 population after cross of *Br prt6.2-12* and *Br prt6.3-1* containing wild-type alleles of *Br PRT6.2* and *Br PRT6.3*. Used as additional WT control for remaining background SNPs caused by EMS mutagenesis. | This study |
| *Br ate1-2* | Contains early stop codon in *Br ATE1* (Bra009127). Obtained from the JIC; line #JI31799-B. | This study |
| *Br ate2-2* | Contains early stop codon in *Br ATE2* (Bra034856). Obtained from the JIC; line #JI31089-B. | This study |
| *Br ate1 ate2/+* | Homozygous mutant for *Br ate1-2* and heterozygous for *Br ate2-2.* | This study |

**Supplementary Table S2: List of oligonucleotides used in this study.**

| **Oligo name** | **Oligonucleotide sequence (5’ -> 3’)** | **Application** |
| --- | --- | --- |
| BM28 | GGTTCTTGGATGGCAAACTAGATGT | Genotype *Br ate1-2* by Sanger sequencing |
| BM29 | GGTAACTTACTTGAAACGAAGAGGACAC | Genotype *Br ate1-2* by Sanger sequencing |
| BM30 | CGTAGTCTCTGAATGATAAACTTACTC | Genotype *Br ate1-2* by Sanger sequencing |
| BM31 | GATCTCTCTCTCAGAAATGAGAACAAG | Genotype *Br ate2-2* by Sanger sequencing |
| BM36 | GCTCCCGATCTCCAGAGAATAAATGTTTCCTC | Genotype *Br prt6.3-1* by Sanger sequencing |
| BM37 | CATTGTATTCTTCCCACAACGAGTCAC | Genotype *Br prt6.3-1* by Sanger sequencing |
| BM93 | CACATATCGTGGATCTTGCACA | Genotype *Br prt6.2-12* by dCAPs |
| BM94 | GCTTCATCGAACCTACCGTT | Genotype Br *prt6.2-12* by dCAPs |
| BM97 | TCTGACGGACATGTAAAGCACTCTTTGaTC | Genotype *Br prt6.2-12* by dCAPs (small-cap letters indicate mismatches to create restriction site) |
| BM98 | CCTCCTGGAACTTCTTTGCAGCC | Genotype *Br prt6.2-12* by dCAPs |
| BM34 | CGATTCCTGTGTCTGGGTCTACTAATG | Genotype *Br prt6.2-12* by Sanger sequencing |
| BM35 | CATCGAATTCTTCGCACAACGAGTAAT | Genotype *Br prt6.2-12* by Sanger sequencing |
| qBM333 | TCTATAGCGTGTCTCTTGTTCAGACTGT | *Br PRT6.1* RT-qPCR |
| qBM334 | CCCAACCACCTGATTCTCTCAAAGC | *Br PRT6.1* RT-qPCR |
| qBM331 | TCTTCTGATACGGCGGACCACAAT | *Br PRT6.2* RT-qPCR |
| qBM332 | CGTTCTTCATTCAGGTAGAGTGGCTT | *Br PRT6.2* RT-qPCR |
| qBM329 | AGGGAGAATTTCTCTGTCCTGCATG | *Br PRT6.3* RT-qPCR |
| qBM330 | GCAGCAGACTGTAGTAGACACAACG | *Br PRT6.3* RT-qPCR |
| qBM234 | TGTCGTTGATGAACACTTTGAGGTGAC | *Br HB1* RT-qPCR |
| qBM235.1 | CGGTGACCACATCTCTGGCAC | *Br HB1* RT-qPCR |
| qBM229 | GGTCGCTGATTCTCCCCAGC | *Br PCO2* RT-qPCR |
| qBM230 | CCGTGAAGTCTGAATCCACTTTCACT | *Br PCO2* RT-qPCR |
| qBM238 | TGCTCAGGCTCAGTTGGTGG | *Br HRE2* RT-qPCR |
| qBM239 | GCCTCTGCCTTATCCCTCTGTAC | *Br HRE2* RT-qPCR |
| qBM225 | GCTTATTGACAGAGTTGCTTGGCACA | *Br MPK3* RT-qPCR |
| qBM226 | CAACAGTGATTCTTTTGCTGGGGTCA | *Br MPK3* RT-qPCR |
| qBM227 | CGGTTAAGATTGTCAAGGTGGCTGT | *Br RBOHD* RT-qPCR |
| qBM228 | CGCTCACGTAGTCGTCTCCTG | *Br RBOHD* RT-qPCR |
| qBM55 | CACCACCGAGTACATGACGTACA | *Br GAPDH* RT-qPCR |
| qBM56 | TGCCCGTGAACACTGTCGTA | *Br GAPDH* RT-qPCR |
| KG59 | GCGGTCGGTAAAGTTGTTCCAT | *LUC* RT-qPCR |
| At327 | GAAGTGTTCGTCTTCGTCCC | *LUC* RT-qPCR |

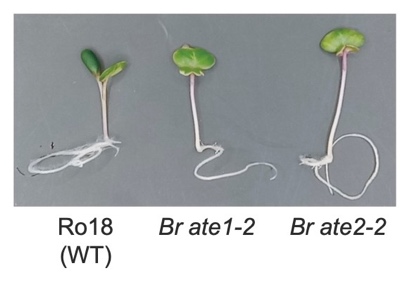

**Figure S1: Single mutant *Br ate1-2* and *Br ate2-2* seedlings are similar to Ro18 wild-type seedlings.** Seeds of each genotype were sown on 0.5x MS agar supplemented with 0.5% sucrose. Seedlings were grown in continuous light for 6 days.

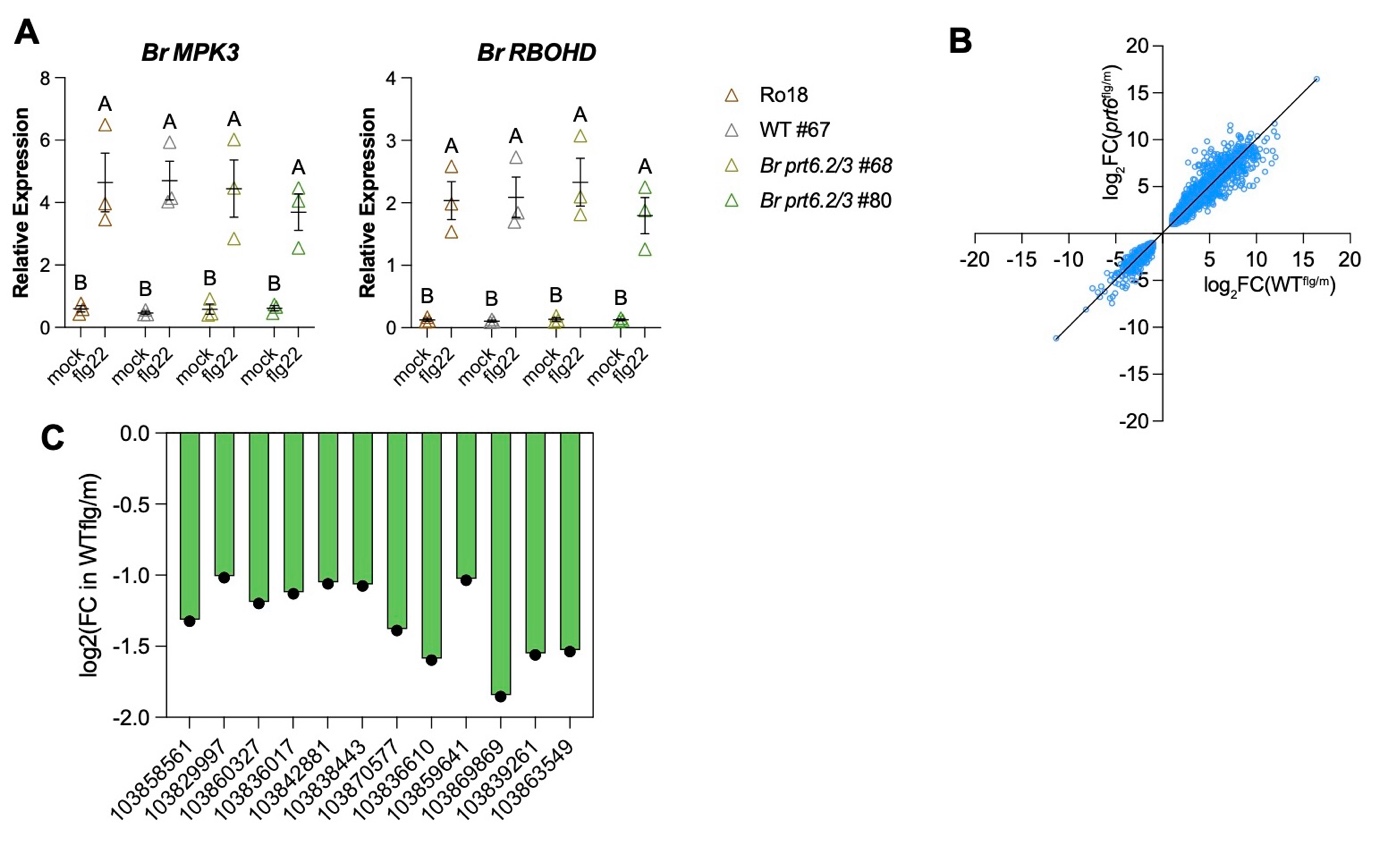

**Figure S2: Supplemental analysis of RNA-seq experiment with flg22. (A)** RT-qPCRs to verify induction of flg22 response genes after 1 hr of treatment of 3-day-old *B. rapa* seedlings of different genotypes. Mean of 3 biological replicates with SEM. Statistical analysis: 2-way ANOVA with Tukey’s test. **(B)** Comparison of directionality and amplitude of gene expression changes amongst DEGs common to wild type and *Br prt6.2/3* mutants in response to flg22. **(C)** Gene expression changes as log_2_(fold change) of DEGs in wild-type seedlings only that belong to the GO categories shown in Figure 4D.

**References**

**Stephenson P, Baker D, Girin T, Perez A, Amoah S, King GJ, Ostergaard L** (2010) A rich TILLING resource for studying gene function in Brassica rapa. BMC Plant Biol **10:** 62
